# Transmembrane domains of H1 and H3 hemagglutinin contribute to membrane fusion in a different manner

**DOI:** 10.1101/2024.10.31.621245

**Authors:** Paulina Borkowska, Szymon Kubiszewski-Jakubiak, Michał Bykowski, Łucja Kowalewska, Michał Michalski, Piotr Setny, Remigiusz Worch

## Abstract

Hemagglutinin (HA), a fusion protein of influenza virus, has been extensively researched as a model fusion viral protein. However, most of the efforts focus on the fusion peptide (FP) and the ectodomain, while relatively little is known about the “membrane anchor”, a transmembrane domain (TMD). While the structural insights on H1 subtype full-length HA underlay the influence of TMD on the ectodomain orientations, the structures representing the other phylogenetic group are still unavailable. Inspired by the sequential varieties of TMDs in both groups, we performed a series of experiments on full-length HA proteins with H1and H3-swapped TMD domains in virus-like particles (VLP). In parallel, we studied the behaviour an and interplay of FP and TMD-corresponding fragments with the use of artificial membrane systems and molecular dynamics simulations. We detect clearly a distinct interplay between FP and TMDs in the two phylogenetic groups. We observe that TMD-dependent fusion activity originates from TMD-lipid interactions, but not direct FP:TMD complexing.

## Introduction

Influenza is a respiratory disease that has affected humans since ancient times (Potter & Jennings, 2011). It currently impacts 10-20% of the global population annually, causing many deaths despite a low mortality rate. A major challenge is its high mutation rate, which increases pandemic risk. The virus spreads through respiratory droplets and enters host cells via endocytosis, with the final step involving fusion of viral and endosomal membranes to release genetic material (White & Whittaker, 2016). The influenza A virus is usually spherical or ovoid, 80-120 nm in diameter (Vajda et al., 2016). Hemagglutinin (HA) and neuraminidase are the most abundant surface proteins. HA is a homotrimeric protein essential for viral entry into host cells, initiating fusion of viral and endosomal membranes (Hamilton et al., 2012). HA has two subunits, resulting from HA0 precursor cleavage: HA1, for binding to cell receptors, and HA2, for the fusion process (Blijleven et al., 2016). The HA subtypes are divided into two main phylogenetic groups based on amino acid sequences and 3D structures of their ectodomains (Lazniewski et al., 2018).

The HA facilitates viral entry into a host cell by first binding to sialic acid on host membrane receptors, initiating endocytosis. When the endosomal pH lowers, HA partially refolds, forming an extended coiled coil stem structure (Benton et al., 2020). HA then inserts fusion domains into the endosomal membrane (Gui et al., 2016). Anchored in the viral membrane through C-terminal transmembrane domains (TMDs) (Benton et al., 2018) and cytoplasmic tails (CTs) (Mineev et al., 2013), and in the endosomal membrane through N-terminal fusion domains (Lorieau et al., 2010; Michalski & Setny, 2022), HA performs a jack-knife motion, bringing the membranes into close contact. This action overcomes the dehydration barrier, leading to a stalk structure, the first fusion intermediate (Fuhrmans et al., 2015; Michalski & Setny, 2023b). During this process, rigid parts of the ectodomain must tilt significantly relative to their membrane anchors.

While structures of the HA ectodomain have been available for several decades (Bizebard et al., 1995; Bullough et al., 1994; Chen et al., 1998, 1999; Eisen et al., 1997; Gamblin et al., 2004; Sauter et al., 1992; Stevens et al., 2004; Weis et al., 1988; Wilson et al., 1981), the first full-length HA structure, comprising the H1 TMD, was published in 2018 (Benton et al., 2018). This full-length structure underlies the interplay between the TMD and the ectodomain conformation. Namely, the TMD was observed at angles between 0° and 52° relative to the ectodomain, indicating a flexible linker region composed of a conserved glycine, a short α-helix, and an extended chain (residues 175–184). In complex with a FISW84 Fab antibody, this angle is reduced to 20° or less (Fig. 1D). The ectodomain base lies horizontally relative to the membrane, with conserved glycine residues facilitating flexibility. Recently, molecular dynamics simulations characterised the transitions between straight and tilted metastable TMD configurations which were supporting the tilting motion of HA ectodomain (Michalski & Setny, 2023a).

**Fig. 1.**
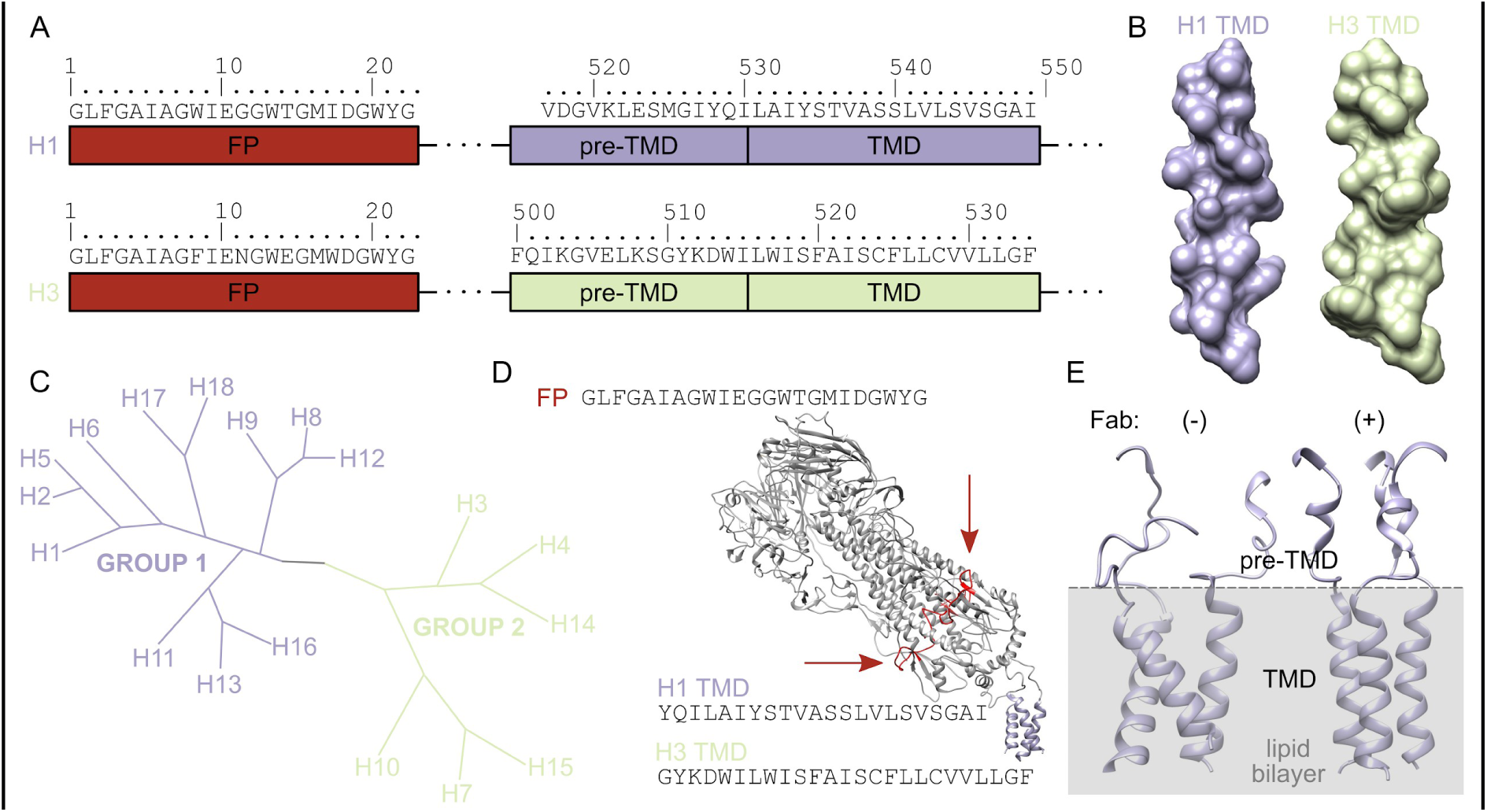
**(A)** Fusion peptide (FP), transmembrane domain (TMD) and pre-TMD sequences of H1 and H3 influenza hemagglutinin (HA) subtypes. **(B)** Surface representation of H1 and H3 TMD built as regular alpha-helices based on sequences. **(C)** Phylogenetic tree of Group 1 and Group 2 hemagglutinins (based on Łaźniewski et al.). **(D)** Sequences of FP, H1 TMD and H3 TMD peptides used in this study and the tilted ectodomain of HA. Fusion peptides (red) marked with arrows. Based on x PDB (Benton et al. 2018). **(E)** Conformation of TMD with the bound and unbound antibody by the ectodomain. All structures rendered with Chimera software (https://www.cgl.ucsf.edu/chimera/).

However, apart from the early studies showing the loss of function of hemagglutinin or parainfluenza virus fusion protein (Kemble et al., 1994; Melikyan et al., 1995; Nüssler et al., 1997; Tong & Compans, 1999) upon replacement with a glycosylphosphatidylinositol (GPI) membrane anchor, little is known about the role of the TMD in membrane fusion. Moreover, there are no experimental data on the structure of H3 TMD, belonging to the other phylogenetic group. As a continuation of our hypothesis on a possible distinct behaviour of HA in membrane which was based solely on physicochemical properties of HA TMDs (Kubiszewski-Jakubiak & Worch, 2020), in this paper we are aiming to fill this gap with the use of experiments. We focus our attention on the TMD-swapped, full-length HA in the form of protein present in plant-derived virus-like particles (VLPs), isolated peptides in artificial membrane systems and molecular dynamics simulations.

## Results and Discussion

### The TMDs from Group 1 and Group 2 behave differently in the membrane bilayer

To assess the possible differences of the H1 and H3 TMD fragments resulting from sequential variations (Fig. 1), we employed full atom molecular dynamics (MD) simulations to characterise the structure of these fragments in lipid bilayer. We performed the MD simulations of TMD-corresponding peptides, further used in experiments, similarly to our protocols applied in fusion peptide studies (Worch et al., 2017, 2018, 2021). Fig 2D shows the histograms of distances between triplets o atoms representing the ectodomain (Leu512:CA or Leu495:CA for H1 and H3, respectively) and TMD (Tyr535:CA or Trp517:CA for H1 and H3, respectively) which may be regarded as an indication of HA ectodomain tilt. We found that the H3 TMD fragment adopts more straightened conformations as compared to the H1 TMD fragment. Surprisingly, this is very much in line with the hypothesis suggested solely based on the basis of the analysis of physico-chemical parameters (Kubiszewski-Jakubiak & Worch, 2020).

**Fig. 2.**
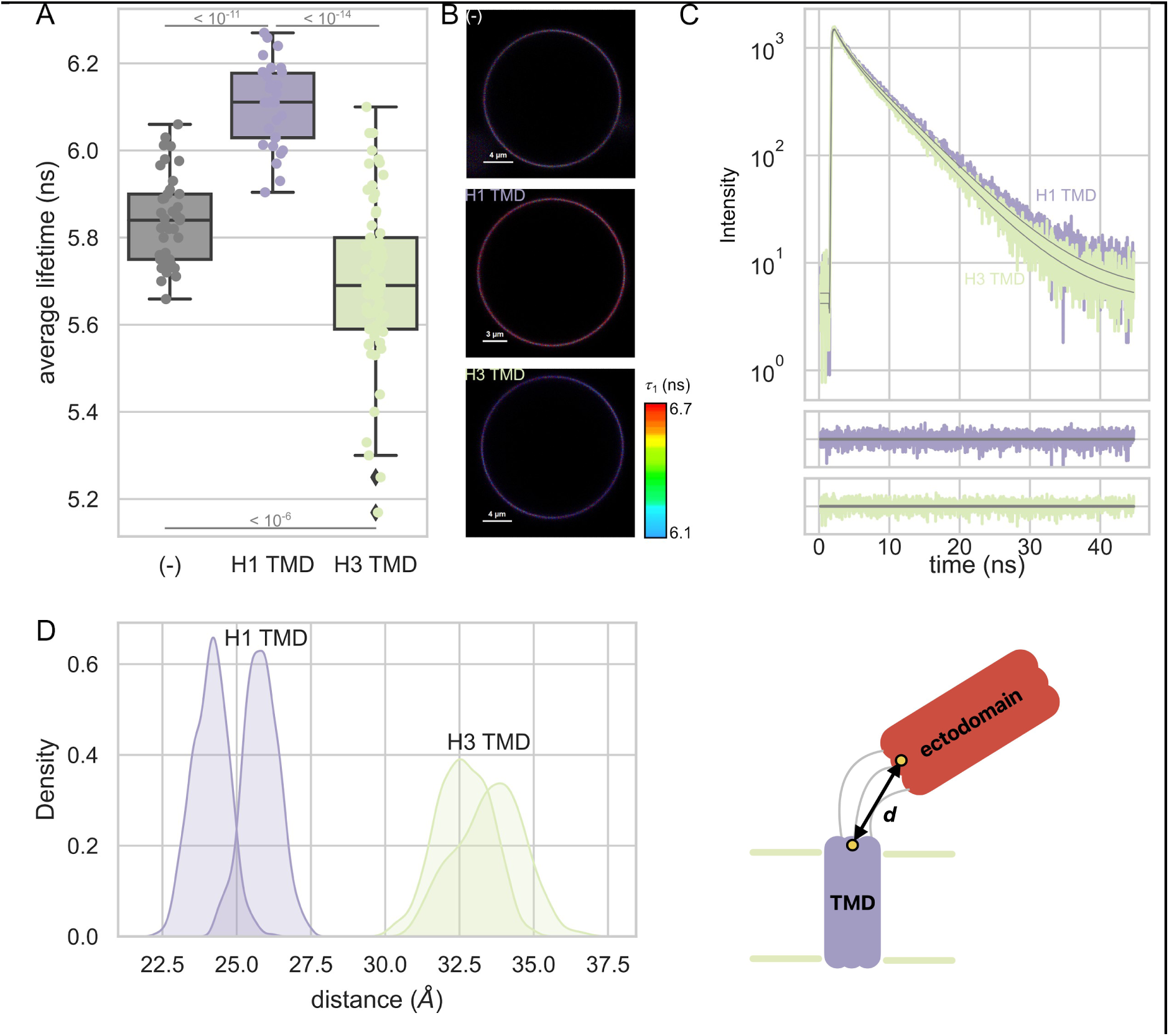
**(A)** Average lifetime values of NBD-C6-DOPE lipid dye sensitive for membrane order with H1 TMD and H3 TMD corresponding peptides incorporated. p-values calculated from Mann-Whitney U test. **(B)** Examples of FLIM images of individual GUVs. **(C)** Example curves of fluorescence decays with double exponential curves fit. **(D)** Histograms of the distances, d, between ectodomain and TMD (see text for description) in the H1 and H3 segments resulting from full-atom MD simulation. **(E)** Pictogram of the distances, d, between ectodomain and TMD in the H1 and H3 segments.

We next evaluated the behaviour of TMD-corresponding peptides in the lipid bilayer experimentally. We utilised the membrane of giant unilamellar vesicles (GUV) composed of phosphatidylcholine (POPC) doped with a fluorescently tail-labelled phospholipid (POPE-NBD-C6), commonly used for determination of membrane order/disorder measurements (Stöckl & Herrmann, 2010). The dye changes its lifetime from low in membrane composed with unsaturated lipids to high when placed in the bilayer exhibiting larger membrane order. To verify the influence of the TMD corresponding peptides on the surrounding lipid tails, we performed a series of FLIM measurements of individual GUVs doped with the TMD peptides (Fig. 2C). To show that the peptides are incorporated into the bilayer, we analysed the fluorescence resonance energy transfer between either tryptophan (H3) or tyrosine (H1) and dansyl labelled lipids in the form of small unilamellar vesicles SUV (SI Fig. 1). Unexpectedly, we found that the presence of TMD changes the order of lipids in a different manner, namely H1 TMD induced a membrane order increase, whereas H3 TMD had the opposite effect. In all cases, the fluorescence decay could be described precisely by a two-component exponential decay with good accuracy (Fig. 2D; more fitting examples are presented in SI Fig. 2).

Membrane ordering or disordering properties of TMD fragments we described above could pave the road towards better understanding of molecular details standing behind the TMD-lipids interactions. H1 TMD, having smaller accessible surface area (ASA) than H3 TMD (Kubiszewski-Jakubiak & Worch, 2020) could fit better between the spaces of unsaturated POPC lipid chains leading to their further disordering. In the case of H3 TMD, in spite of its larger bulkiness, in general decreasing the raft affinity (Lorent et al., 2017), the presence of cholesterol consensus motif (CCM) (Hu et al., 2019) might facilitate the ordering of H3 TMD segment next to large and flat surfaces of cholesterol rings. Ordering effect of the TMD belonging to Group 2 was also observed by means of electron spin resonance (Ge & Freed, 2011). The aforementioned observations open a question whether a diverse effect of the TMD segments on lipids itself could be translated to HA fusion functions.

### Full-length hemagglutinin in virus-like particles (VLPs)

It has been shown that mutations of the CCM YKLW motif in the TMD of H3 did not disturb the transportations of virons to the apical membrane of polarized cells, however had an effect on its fusion proteins measured in cellular assays (Hu et al., 2019). We asked a question whether an observed, distinct behaviour of the TMD segment in the bilayer has any effect on the functional properties of hemagglutinin. To answer that, we performed a series of experiments with the use of plant-derived virus-like particles (VLP), which allowed for studying the full-length HA embedded in lipid bilayer. The other benefit of VLPs is that HA is organised in a similar fashion as in influenza viron. Since we intended to study the influence of the TMD segments, we prepared two versions of the plasmid, the native full-length H1, representing Group 1 (abbreviated further as H1-TMD-H1) and H1 with the swapped TMD and pre-TMD fragments from the H3 subtype (H1-TMD-H3), representing Group 2 (Fig. 1C). Since the major sequential differences are found in the TMD and pre-TMD regions and the fusion peptide sequences are very similar (Langley et al., 2009; Worch, 2014), preparation of the reverse exchange (i. e. H3-TMD-H1) was not necessary. Regarding the VLPs preparation steps, in short, the HA-encoding plasmids were introduced to *Agrobacterium* from which they were further used in the transformation of *N. Benthamiana* plants (around 4 weeks old) (see Materials & Methods for details). Hemagglutinin expressed in such a system was previously shown to be secreted in vesicles located between plasma membrane and cell wall in the leaf tissue (D’Aoust et al., 2008, 2010; Peyret, 2015), similarly as in our experiments (Fig. 3A). More of the TEM pictures of leaf fragments, including not-transformed leaves pictures as controls, are present in SI Fig 3. We could confirm the presence of hemagglutinin in harvested VLPs by Western blot (Fig. 3B) which also showed that the majority of the protein is expressed in its uncleaved, precursor (HA0) form. To analyse the size of the harvested VLP (see Materials and Methods for details), we measured the dynamic light scattering (DLS), which resulted in monomodal distributions of sizes with the averages of 142 nm and 145 nm, for H1-TMD-H1 and H1-TMD-H3, respectively (Fig. 3C). Thus, our method of choice resulted in obtaining virus-like particles with average sizes very close to the complete, natural viron particles.

**Fig. 3.**
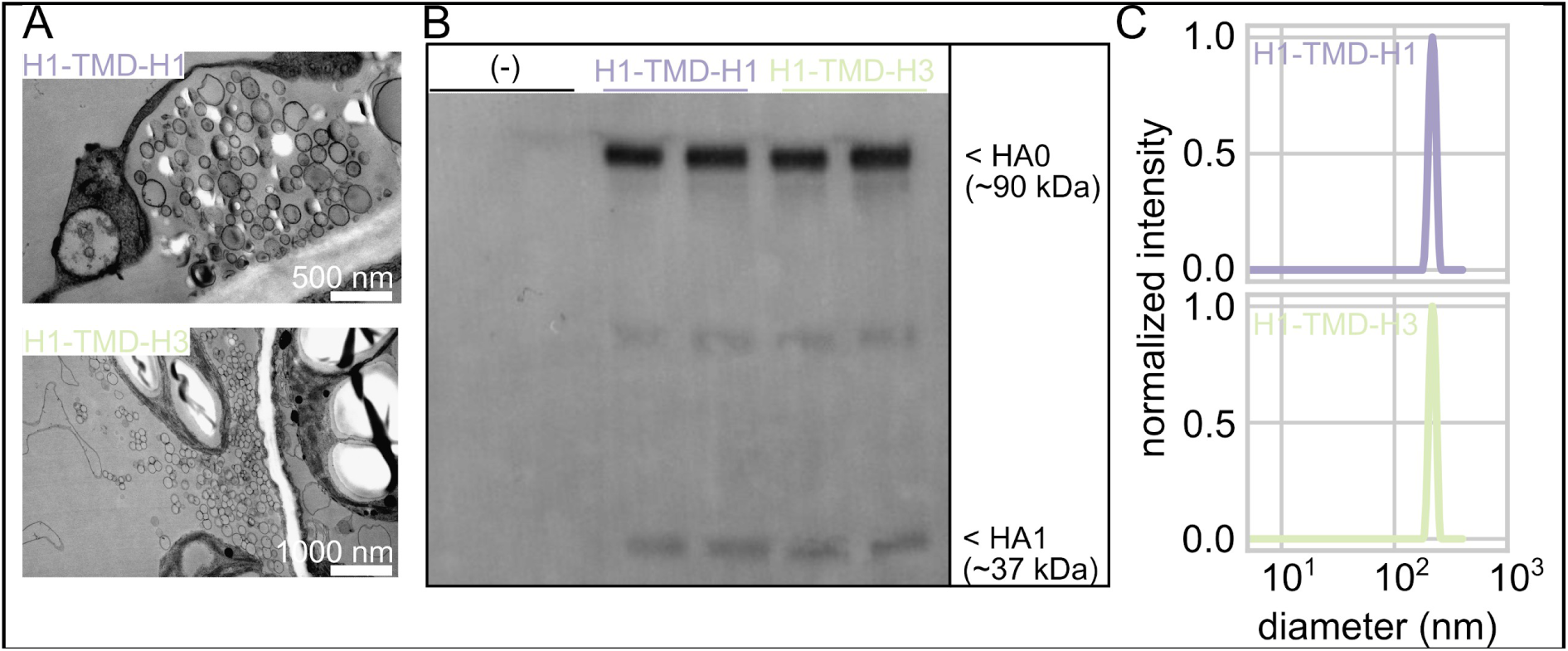
**(A)** Transmission electron microscopy pictures of leaves fragments from transformed *Nicotiana benthamiana* plants with the H1-TMD-H1 and H1-TMD-H3 containing virus-like particles. **(B)** Western blot prepared from harvested VLPs (two repetitions each). **(C)** Histograms of diameters of harvested VLPs obtained from dynamic light scattering (DLS).

### Fusion efficiency is TMD dependent

To corroborate the fusion activity of HA present in plant-derived VLPs, we first investigated the fusion efficiency of VLP and other lipid membrane in the form of a cushioned-supported lipid bilayer (SLB) prepared on glass (Fig. 4 C). For this purpose we optimised the preparation of a SLB with the dope of 5 mol% DOPE-polyethylene glycol (DOPE-PEG 5k) in the lipid mixture. The presence of PEG moieties ensured the distance between the glass support and the membrane, resulting in the preservation of lateral mobility of lipids, as corroborated by fluorescence correlation spectroscopy (FCS) (SI Fig. 4). Next, we designed the experiment in the following way: we introduced the donor (NBD-PE) in the supported lipid bilayer in order to observe the fluorescence resonance energy transfer (FRET) which might occur if the molecules of the acceptor (N-Rh-PE) are in close proximity. To quantify FRET efficiency, we performed a series of FLIM images in order to determine the fluorescence lifetime of a donor which should decrease in the case of occurring FRET. To test this approach, we started with the positive and negative controls. In the former one, both dyes are placed together in the mixture whereas in the latter case, 100 nm LUVs doped with the acceptor were added and incubated as the supernatant. Under such conditions, we observed changes in fluorescence lifetime of the donor; it did not change after LUV incubation, however, it decreased in the positive control (Fig. 4A). This result also indicated that both fluorescently labelled lipids must be present in the same bilayer in order to observe the FRET effect. In other words, it is not possible to detect FRET between the donor located in the SLB and the acceptor present in the unfused liposome. Once we had established the conditions of control experiments, we performed the experiments with N-Rh-PE-labelled VLP. Incubation of activated, proteolytically cleaved H1-TMD-H3 led to a larger decrease of the mean fluorescence lifetime in the donor, as compared to the parallel experiment with the H1-TMD-H1. Together, these observations show first of all that the HA present in plant-harvested VLP preserves its activity and, secondary, the presence of the H3 TMD segment intensifies HA fusion abilities.

**Fig. 4.**
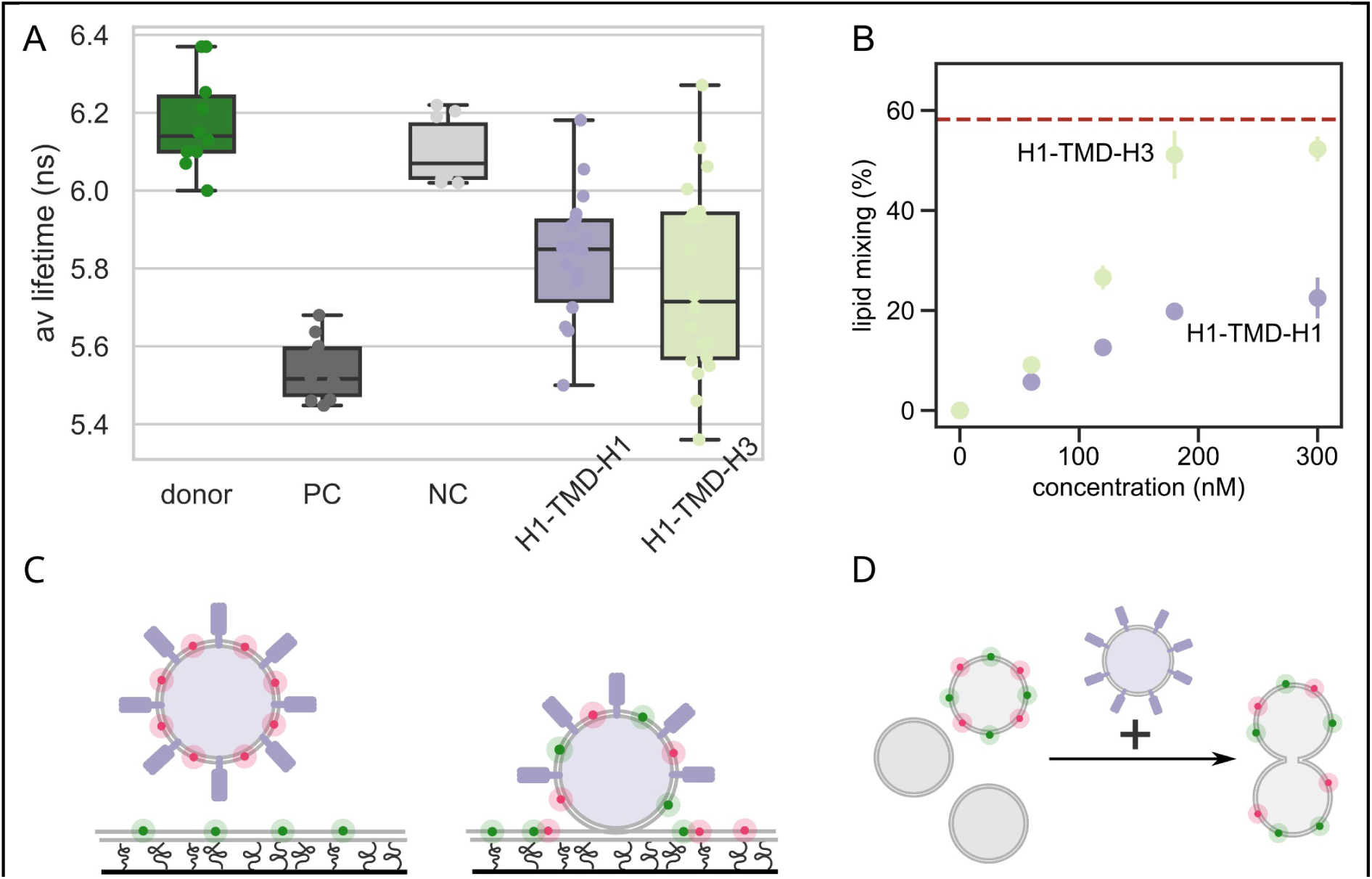
**(A)** Average lifetime values of the donor placed in a supported lipid bilayer. Positive and negative controls (PC and NC, respectively) for FRET with the acceptor present in the same bilayer (PC) or present in the acceptor labelled LUVs (NC). Lowered values of lifetime for incubated acceptor-labelled H1-TMD-H1 and H1-TMD-H3 VLPs (p-values for H1-TMD-H1 and H1-TMD-H3 mutually with other data <10^−5^. **(B)** Lipid mixing between labelled and unlabelled pools of LUVs fused by hemagglutinin (HA) present in H1-TMD-H1 or H1-TMD-H3 LUVs as a function of HA concentration. Dashed red line corresponds to the fusion activity reached by isolated fusion peptide. **(C)** A scheme showing the FRET experiment between the donor-containing cushioned SLB and VLP-donor doped acceptors. **(D)** VLPs acting as a fusion agent in lipid mixing experiment.

Next, to extend the studies of VLP-induced membrane fusion, we performed the lipid mixing experiments in bulk volume (Fig. 4B). Here, the main focus was directed on the ability of inducing the fusion between the liposomes of the same kind rather than the fusion of the VLP lipid envelope itself. Using such an approach, the VLP pools are serving as fusing agents and performing such an experiment in a bulk volume had a benefit of a better control of HA amount. Fig 4B shows the results of lipid mixing efficiency as a function of HA concentration. To our surprise, in native plant membrane composition H1-TMD-H3 outperformed H1-TMD-H1 with the plateau reaching the lipid mixing levels obtained by an isolated fusion peptide (FP) at a saturating concentration). This result shows an interplay between TMD which, thanks to the interaction with surrounding lipids leads to efficient exposition of FP. Notably, all fusion experiments were performed at pH5, as in matured endosomes, after proteolytic cleavage. As a control, we carried out similar experiments at pH 7 with activated VLPs and observed almost no fusion activity (SI Fig. 5). This shows preservation of native HA behaviour in VLPs.

Obviously the composition of the lipid envelope of plant-derived VLPs varies from an envelope built out of animal plasma membrane. Although cholesterol does not occur in plant membranes, there is still a variety of other sterols which are quite abundant (Cacas et al., 2016; Suza & Chappell, 2016). It is known that the virus envelope is cholesterol-enriched and probably ‘raft-like’ environment is important for functioning of hemagglutinin. Since the notions of raft-like behaviours of plant membranes are more and more often, in the light of membrane heterogeneity based on the coexistence of liquid ordered and liquid disordered phases, there might be in fact many similarities.

Since the H3 TMD - containing structure still remains unsolved, we can only speculate about the possible structural rearrangements which potentially make the fusion peptide more expanded to the target membrane and therefore facilitating eventual fusion. It would be also in line with the results of TMD domain swapping, where H9 TMD (Group 1) was replaced by H3 TMD (Zhang et al., 2017). Such a recombinant attenuated virus showed enhanced immunogenicity in immunised mice and chicken. To underline further differences between the two TMD representatives of Group 1 and 2, we performed a series of experiments aiming at reconstitution of full-length HA in LUVs of desired composition. In short, affinity-purified HA (SI Fig. 6) was mixed with LUVs (compositions: POPC/cholesterol 6/4 % and POPC/sphingomyelin/cholesterol 2/2/21 mol%) in the presence of detergents which were eventually removed by dialysis. HA in such form preserved its activity, assessed by lipid mixing assays (SI Fig. 7). Stunningly, H3-TMD-H1 version of HA also outperformed H1-TMD-H1 when reconstituted in POPC/cholesterol 6/4 mol % membrane. Fusion activity of H1-TMD-H1 was however rejuvenated for the ‘raft’-like membrane composition (SI Fig. 7). Although it is impossible to compare directly the fusion efficiency and MD simulations of the TMD fragments, we observed a distinct behaviour between H1 and H3 TMD in membranes with varying compositions (Fig. 6). In general, in the case of H3 TMD, the distance defined as in Fig. 2 E, spans a wider range as compared to H1 (Fig. 6). Probably a more versatile, membrane-adaptable positioning of H3 TMD is translated into structural rearrangements leading to more efficient fusion. Understanding the principles standing behind coupling of the lipid-TMD interactions and HA fusion properties and antigen presentation could contribute to the development of novel antiviral drugs and efficient subtype-specific vaccines.

### Fusion activity is not a result of direct FP:TMD interactions

To verify the possibility of direct, synergistic interactions between FP and TMD we have performed a series of experiments with the use of isolated peptides corresponding to the FP, H1 TMD and H3 TMD sequences (Fig 1. D). As a reference we used the FP, known to have fusogenic properties by itself and to cause pore formation in the membrane leading to efflux of encapsulated fluorescent dye (Chakraborty et al., 2013). For this purpose, we measured the lipid mixing and leakage efficiency for isolated peptides as well as for FP:TMD mixes. In none of the cases the presence of the TMD fragment did not show a significant membrane activity and did not cause an increase of FP activity (Fig. 5). Moreover, the effects were rather contrary (*i. e.* the presence of TMD lowers the activity of the FP) and were more pronounced in the case of H1 TMD for the liposomes containing cholesterol. This part of results also facilitates the interpretation of the experimental results with the application of the full-length HA, since it shows the lack of interplay between FP and TMD hemagglutinin parts. This issue is still controversial in the literature, since interaction between FP and TMD were pointed as important driving factors on the way to the fusion pore formation by means of e.g. electron spin resonance (Lai & Freed, 2015) and fluorescence and electrophoresis (Chang et al., 2008). However in a hydrogen-deuterium exchange mass spectrometry (HDX-MS) on the complete HA2 subunit FP:TMD complex was not supported. In our opinion, care must be taken when making the “zero-one” type conclusion, since the interpretation may depend on a number of factors, which are often even omitted in the experimental descriptions. For example, the termini form of the peptides used. Our results point rather the involvement of a TMD segment in the activity of an entire “fusion machine” which serves a way more than a passive membrane zipper. A tremendous progress in structural biology taking place in recent years hopefully will help to answer these burning questions.

**Fig. 5.**
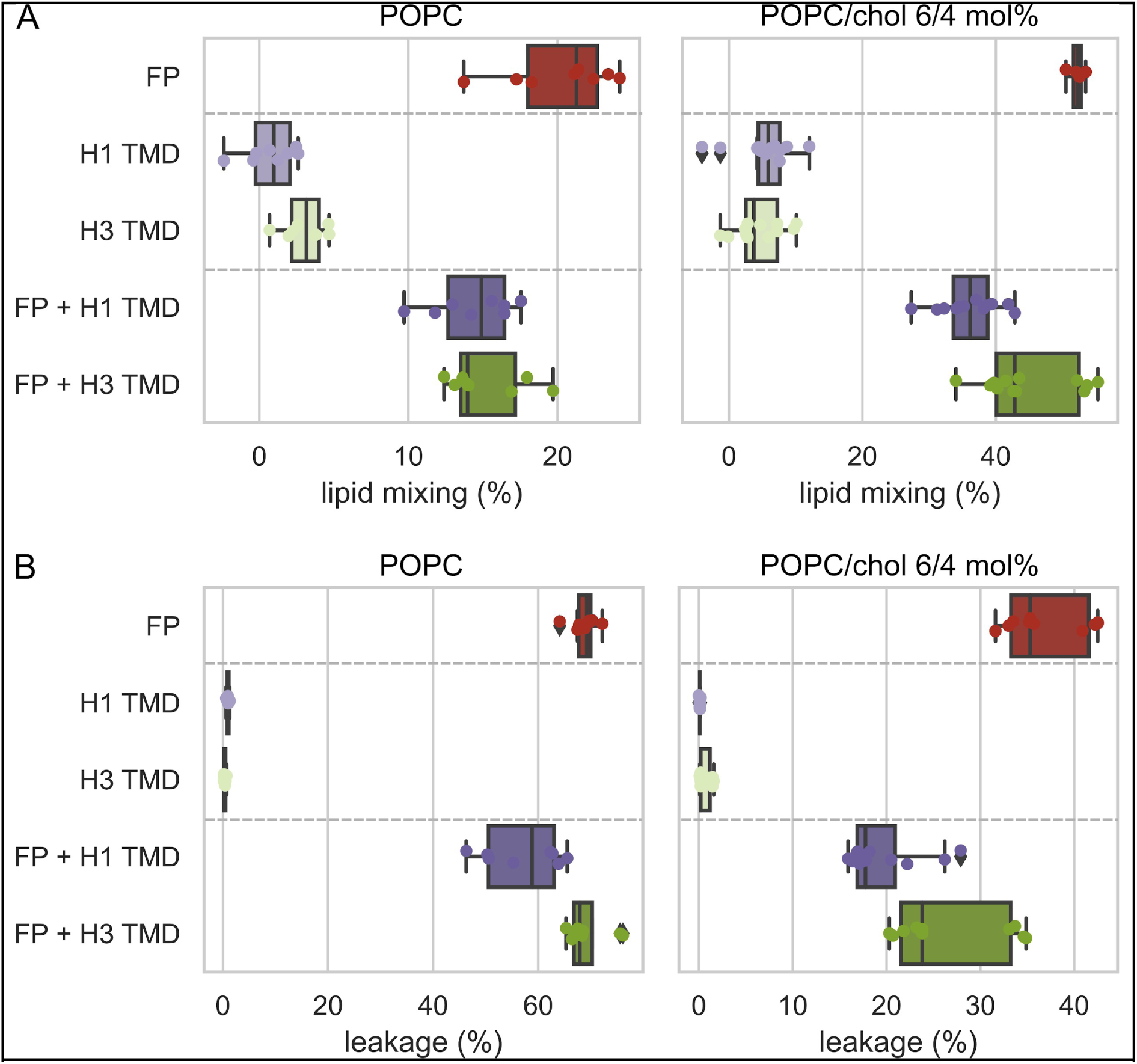
Membrane activity of FP and H1 TMD or H3 TMD isolated peptides and their mixes measured as **(A)** lipid mixing and **(B)** leakage of encapsulated fluorescent dye. Two lipid compositions were compared: pure POPC and POPC/cholesterol 6/4 mol% mixture (left and right panels,respectively).

**Fig 6.**
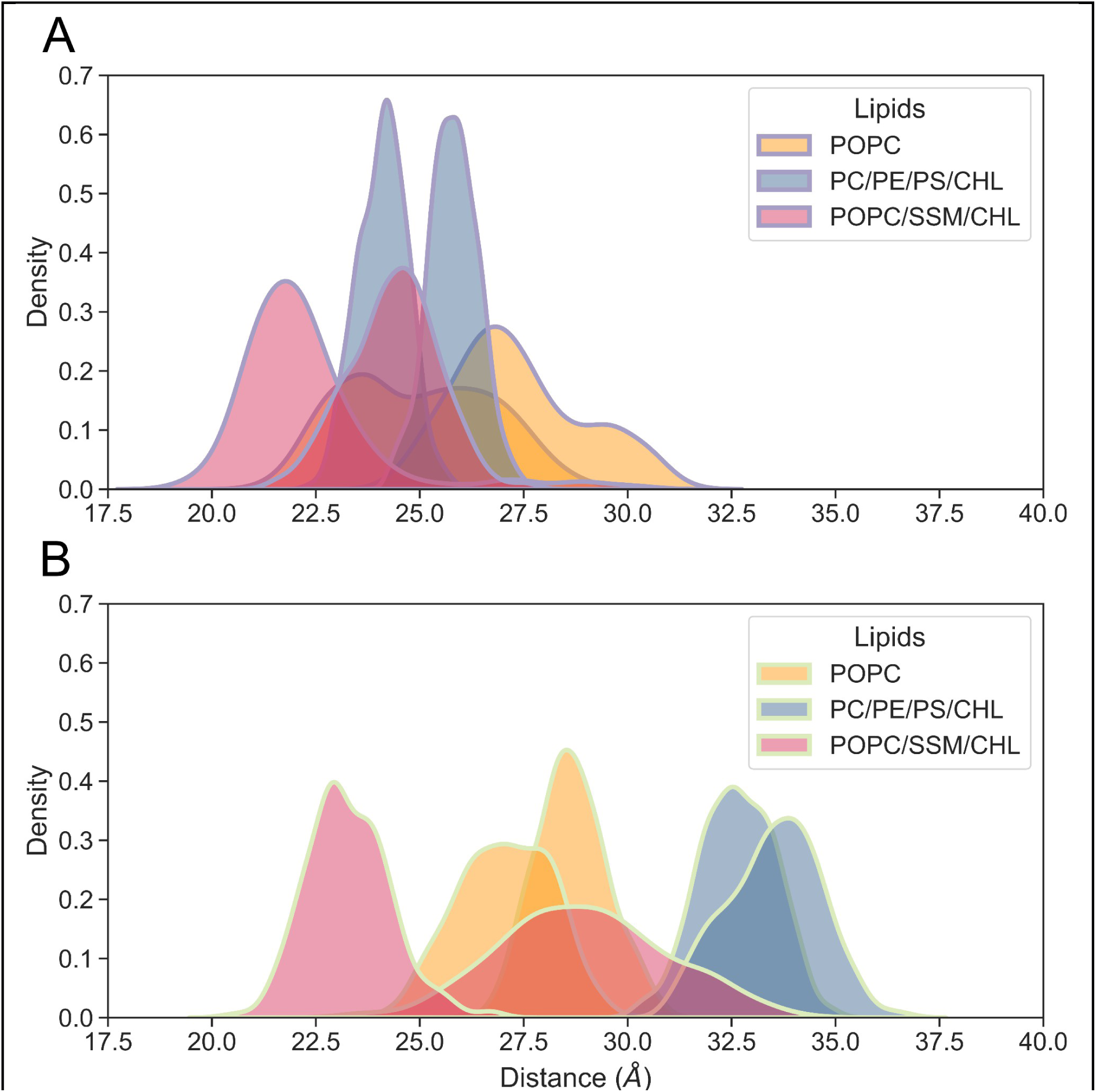
Histograms of the distances in the TMD segments resulting from full-atom MD simulation of the different lipids composition. **(A)** MD simulation for the H1 segment, **(B)** MD simulation for the H3 segment.

## Conclusions

Here we analyse, the effect of the transmembrane domain type on the surrounding lipids and the fusion function of the full-length hemagglutinin. Studies of a full-length membrane protein were facilitated by the application of plant-derived virus-like particles which, naturally formed as a result of transformation, could serve as a functional platform for quantitative experiments. This study also implies that concluding about the protein behaviour without the membrane context does not provide in-depth understanding. Indeed, more examples of coupling the proteins and lipids are reported as regards viral membrane fusion (reviewed in White *et al*. 2023). Studying the bridging of the transmembrane domain influence of activity and the antigen presentation by viral fusion proteins calls for further investigations and opens up a plethora of potential applications.

## Acknowledgements

The research was funded by the Sonata Bis 8 grant (National Science Centre UMO-2018/30/E/NZ1/00257). TEM studies were performed in the Laboratory of Electron Microscopy of the Nencki Institute, supported by the project financed by the Minister of Education and Science based on contract No 2022/WK/05 (Polish Euro-BioImaging Node “Advanced Light Microscopy Node Poland”).

## Materials and Methods

### MD simulations

MD simulations were used to analyze the tilting of H1 and H3 ectodomain within lipid membrane environment. Initial ectodomain structure was modeled based on cryo-electron microscopy (cryo-EM) data for H1, with PDB ID 6HJQ. H3-specific point mutations, missing protein regions, including the N-terminus representing the ectodomain and linker, as well as the C-terminus of the TMD, were reconstructed using BIOVIA Discovery Studio 2021. For the C-terminus, we assumed an α-helical conformation for the entire TMD region and an extended structure for the CT fragment to avoid any preliminary bias. The structures also included post-translational palmitoylation of cysteine residues within the CT fragments.

Each construct was embedded in four distinct membrane environments: (a) a viral-like membrane composed of 13.3% 1-palmitoyl-2-oleoyl-glycero-3-phosphocholine (POPC), 23.3% 1-palmitoyl-2-oleoyl-sn-glycero-3-phosphoethanolamine (POPE), 18.3% 1-palmitoyl-2-oleoyl-sn-glycero-3-phospho-L-serine (POPS), and 45.0% cholesterol (CHL); (b) a pure POPC (100%) lipid bilayer; (c) a mixture of 55% POPC and 45% CHL; and (d) a membrane consisting of 33.3% POPC, 33.3% SSM, and 33.3% CHL. The protein-membrane systems were immersed in an aqueous solution containing 150 mmol/L Na^+^ and Cl^-^ ions, along with additional ions to neutralize the system’s overall charge. The systems were built using the CHARMM-GUI server (Jo et al., 2008). All molecules were parameterized using the CHARMM36M force field (Huang et al., 2016), and water was modeled with the TIP3P model. Throughout the simulations, covalent bonds involving hydrogen atoms were constrained using the LINCS algorithm, and a time step of 2 fs was employed. Electrostatic interactions were calculated using the particle mesh Ewald method, with a Lennard-Jones potential cutoff of 1.2 nm and a force-switch smoothing function applied between 1.0 and 1.2 nm. Simulations were conducted at 310 K, controlled by the Nosé–Hoover thermostat, and at 1 bar pressure, maintained by a semi-isotropic Parrinello–Rahman barostat. All MD runs were performed using GROMACS software (Abraham et al., 2015), following the standard CHARMM-GUI protocol for system equilibration. To confirm the stability of experimentally derived TMD configurations and assess the tilting motion of the ectodomain, two 1 μs unrestrained MD production runs were conducted for each studied membrane-bound protein. The analysis was based on the last 500 ns of each run.

### Peptides and chemicals

H1, H3 peptides were custom synthesized by Lipopharm and Biomatik. FP were synthesized by Lipopharm. All phospholipids and cholesterol were purchased from the Avanti Polar Lipids. Experiments were carried out in buffer pH 5 (10 mM citric acid, 150 mM NaCl).

Concentrations of peptides were checked by UV spectroscopy using extinction coefficient at 280 nm of respective peptides. (N-(7-Nitrobenz-2-Oxa-1,3-Diazol-4-yl)-1,2-Dihexadecanoyl-sn-Glycero-3-Phosphoethanolmine (NBD-PE) and 1,2-Dihexadecanoyl-sn-Glycero-3-Phosphoethanolamine Lissamine Rhodamine B (N-Rh-PE) used in fusion assays were from ThermoFisher Scientific. Bis[N,N−bis(carboxymethyl)aminomethyl]fluorescein (calcein) used in leakage assay was purchased from Sigma. 1,2-dioleoyl-sn-glycero-3-phosphoethanolamine-N-(5-dimethylamino-1-naphthalenesulfonyl) (ammonium salt) 18:1 Dansyl PE was purchased from Avanti Polar Lipids.

1,2-dioleoyl-sn-glycero-3-phosphoethanolamine-N-(7-nitro-2-1,3-benzoxadiazol-4-yl) (ammonium salt) 18:1 (C6 NBD PE) used in FLIM was supplied from Avanti Polar Lipids. Acrylamide and Triton X-100 were from Sigma.

### Liposomes preparation

Using standard procedures of electroformation (Angelova & Dimitrov, 1986), Pt wire electrodes were used in the homemade teflon chamber to prepare GUVs. Beforehand each wire was cleaned in ethanol, then we applied 5 μl of 1 mg/ml of the lipid mixture in chloroform, followed by 15 min of drying under vacuum. Then the teflon chamber was filled with 350 μl of 0.3 M sucrose and an AC-current of 3 V (peak-to-peak) was applied in two steps: 10 Hz for 2 h and 2 Hz for 0.5 h. Eventually 40 μl of GUVs solution was transferred to a homemade chamber consisting of cut 1.5 ml Eppendorf tubes glued (Norland Optical Adhesive 63) to a glass cover slip (24 × 60 mm, 1.5H, Carl Roth). Each chamber contained 250 μl of buffer.

LUVs were prepared by drying desired concentration of lipids in chloroform under the steam of the nitrogen and subsequently for 4 h under vacuum. Then we rehydrated it for 2 h by shaking with the desired volume of the buffer to the final lipids’ concentration of 5 mg/ml. In the further preparation liposomes were frozen in the liquid nitrogen and then thawed at a 55°C water bath, repeating this process five times. That followed extrusion through 100 nm polycarbonate filters (Whatman) using Avanti Mini Extruder. The size of the LUVs were controlled by dynamic light scattering (DLS) measurements.

SUVs were prepared by drying desired concentration of lipids in chloroform under the steam of the nitrogen and 4 h under vacuum. Then we rehydrated it for 2 h by shaking with the desired volume of the buffer to the final lipids’ concentration of 5 mg/ml. In the following stages of preparation liposomes were sonicated for 10 minutes using Branson S250D tip sonicator in a cycle: 5 s sonication/5 s pause with 10% amplitude. The size of the SUVs were controlled by DLS measurements.

### Preparation of supported lipid bilayer (SLB)

SLB were prepared from the LUVs with final lipids concentration of 5 mg/ml. For the FCS measurements POPC/chol/PEG-PE/Atto 488 (55/40/5/0.02 mol %) mixtures were prepared. LUVs were diluted by mixing 20 μl of LUV with 130 μl of buffer pH 5 with calcium (10 mM citric acid, 150 mM NaCl, 2 mM CaCl_2_). Then LUVs were placed into a homemade chamber consisting of cut 1.5 ml Eppendorf tubes glued (Norland Optical Adhesive 63) to a glass cover slip (24 × 60 mm, 1.5H, Carl Roth). Glass slides were cleaned in chromic acid for 30 minutes prior to use. The sample was incubated with glass cover slip for 30 minutes. After incubation, SLB was washed 20 times with 200 μl of buffer pH 5 without calcium. Mobility of lipids were checked by FCS measurements.

### Lipid mixing

Lipid mixing enables the measurement of the fusion activity of peptides and VLPs. Experiments were performed on the TECAN plate reader on the Greiner plates (96 well, F-bottom (chimney well) and measured by FRET at room temperature. In these experiments, two sets of liposomes were prepared unlabeled and labeled. Labeled liposomes were prepared by adding 1 mol% NBD-PE and N-Rh-PE to the lipid mixture before drying lipids in the liposome preparation process. Unlabeled and labeled LUVs were mixed in a 9:1 ratio at the final concentration of 100 μM. Fluorescence intensity of acceptor with excitation at 463 nm and emission at 590 nm was recorded. After the initial mixing fluorescence intensity was measured. Subsequently peptides were added to solution in the final concentration of 2 μM and directly after mixing the fluorescence signal was recorded. VLPs were added based on protein concentration, with final concentration of 60, 120, 180 and 300 nM.

Then to verify a 0% fluorescence signal, liposomes were disrupted by adding 4 μl of 10% triton X-100 to the solution and the lowest signal was measured. The experiments were conducted in two independent batches, with separate liposome preparations. The sample sizes were n=12-15 for each VLPs and n=8 for each peptide and mixtures.

### Leakage assay

Leakage assay enables the examination of liposome disruption by membrane-active peptides. The assay was performed using TECAN plate reader on the Greiner plates (96 well, F-bottom (chimney well)) at room temperature. Firstly, calcein was encapsulated at the self-quenching concentration (112 mM of calcein in 0.27M NaOH) in LUVs. The encapsulation involved hydrating a lipid film with a dye solution. In the next step the size exclusion chromatography (PD-10 columns, GE Healthcare) was performed to separate the liposomes and free dye to the fractions. To elucidation a pH 5 buffer was used. The liposome-containing fractions were utilized in further steps. Final concentrations of lipids in the wells were 100 μM. After measuring the initial fluorescence signal (0%), peptides were added to solution in the final concentration of 2 μM. Finally liposomes were disrupted by adding 4 μl of 10% triton X-100 and fluorescent signal representing 100% of leakage was recorded. The fluorescence intensity of calcein with the single excitation/emission wavelengths (495/515 nm) was carried out. The experiment was conducted in two independent batches, with separate liposome preparations. Measurements within each batch were performed in quadruplicate.

### DLS/MADLS Dynamic light scattering/Multi-Angle Dynamic Light Scattering

DLS and MADLS were performed on the Zetasizer Ultra (Malvern). DLS was used to measure size distribution of liposomes and polydispersity of the samples. MADLS enables us to measure the number of particles in the solution (particles/ml). Each measurement was performed with 70 μl of a sample at 25°C.

### Membrane binding of peptides assay

To examine the incorporation of peptides, a membrane binding assay based on FRET was carried out. SUVs were labeled by adding 1 mol % Dansyl-PE to the lipid solution before drying lipids in the liposome preparation procedure (POPC/Dansyl-PE). At the same time non-labeled SUVs were prepared. Dansyl-PE was incorporated into liposomes as the FRET acceptor, while Trp from H3 and Tyr from H1 acted as donors. The assay was performed on the Horiba FluoroMax-4 spectrofluorometer at 20 °C, using cuvette 4 × 10 mm (Hellma). Fluorescence of Trp/Tyr was excited by 280/295 nm, respectively. Fluorescence emission of the dansyl-PE was observed in the range of 420-560 nm for Trp and 420-540 nm for Tyr. The final concentration of lipids and peptides were 2 μM and 100 μM, respectively. Fluorescence was normalized after subtracting background (fluorescence intensity from non-labeled SUVs). The experiment was conducted in two independent batches, with separate liposome preparations.

### FLIM

At first, GUVs with POPC and 0.25 mol% NBD-PE were prepared. Then, the respective peptide was added to the solution containing dyed liposomes and incubated for 30 minutes, followed by imaging. For each peptide ∼ 30 GUVs from the 2-3 electroformation runs were measured. All measurements were performed on Leica TCS SP8 and 63x water-immersion objective (NA=1.2) at RT and 512×512 pixel images were collected. White light laser with a wavelength of 480 nm were used as an excitation of the fluorescence dye with emission spectrum 500-600 nm emission was recorded with the bandpass filter cube 500-550. FLIM images were acquired and analyzed by SymphoTime64 To obtain a fitted curve, the deconvolution method with calculated IRF was performed. Fluorescence decays were fitted with two potential decay equation:

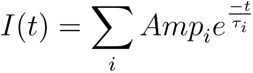

where I(t) intensity in time, amp_i_ amplitude of the i component, _T i_ lifetime of the i component, i= 2.

Average amplitude lifetimes (_Tamp_) from the every GUVs were calculated according to the equation:

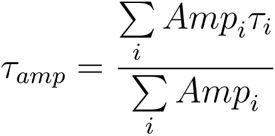

The sample images were pseudocolor-colored corresponding to the longer lifetime (tau1) of the pixels. To acquire fluorescence lifetime, decay curves were fitted as a sum of the exponential terms. Two exponential fitting were sufficient for all GUVs measurements. Examples of fitting and residuals with the Χ^2^ value are presented in SI.

### FCS (Fluorescence Correlation Spectroscopy)

To measure the mobility of lipids in SLB membranes, the FCS measurements were performed. FCS was carried out on Zeiss LSM780 microscope using a 40x water-immersion objective (NA =1.2) at room temperature. For each experiment, the focal volume was calibrated with a 20 nM Alexa488 dye (ThermoFisher) with a diffusion coefficient of 414 μm^2^s^−1^. For fit the fluorescence intensity data for three-dimensional diffusion, the following autocorrelation function, which includes triplet state of fluorophore, was used:

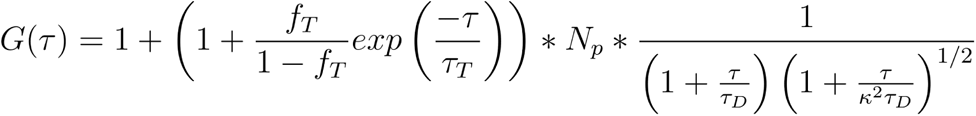

where _T_ is lag time, f is the fraction of fluorescence dye in triplet state, _T_ T fluorescence lifetime of triplet state, N_p_ is a average number of fluorescent particle in focal volume, _T D_ is the diffusion time of the particle, 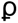 is the structural parameter 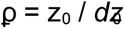 (where *z*_0_ is axial radius and *dz*_0_ is lateral radius of the confocal volume).

Based on obtained *dz*_0_ from fitting equation, the lateral axial radius (*dz*_0_) was calculated:

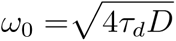

where D is diffusion coefficient

For fitting two-dimensional diffusion in the SLB, the reduced autocorrelation function was used:

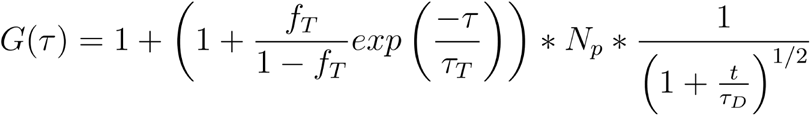

Based on the fitting equation and obtained _T D_ the diffusion coefficients were calculated. The experiment was conducted in two independent batches, with separate SLB preparations.

### FLIM-FRET

To examine the fusion activity of VLPs on the SLB membrane, FLIM-FRET experiment was carried out. FLIM-FRET measurements were performed on Leica TCS SP8 with 63x water-immersion objective (NA=1.2) at room temperature, 512×512 pixel images were collected. FLIM was utilized to examine FRET between NBD-PE (donor) and Rh-PE (acceptor). White light laser with a wavelength of 480 nm were used as an excitation of the NBD-PE with emission spectrum of 490-560 nm, emission was recorded with the bandpass filter 500-550. In the experiment, membrane sections with dimensions of 50×50 μm to a photon count of about 1500 were acquired. For the FLIM-FRET experiment, SLBs with different compositions were prepared using POPC/chol/PEG-PE (55/40/5 mol %) with various dyes composition. Additionally, LUVs with Rh-PE were used as not-fusing negative control. All prepared samples are summarized in Table 1. To examine changes in the donor lifetime, both positive control (PC) and negative controls (Donor & NC) were prepared. Before the FLIM-FRET experiments 100 μl of VLPs were digested with 0.1 mg/ml Xa factor protease (Biolab) overnight at 4°C. They were then dyed with Rh-PE for 2 h at final concentration of 5 μM at RT. VLPs and LUVs were incubated with SLB for 30 minutes before imaging.

**Table 1.** All samples prepared for FLIM-FRET experiments.

| Short name | Composition |
| --- | --- |
| Donor | SLB: POPC/cho/PEG-PE/NBD-PE (55/40/5/0.05 mol %) |
| PC | SLB: POPC/cho/PEG-PE/Rh-PE/NBD-PE (55/40/5/0.25/0.05 mol %) |
| NC | SLB: POPC/cho/PEG-PE/NBD-PE (55/40/5/0.05 mol %)<br>+ LUV: POPC/Rh-PE (100/0.25 mol %) |
| H1-TMD-H1 | SLB: POPC/cho/PEG-PE/NBD-PE(55/40/5/0.05 mol %)<br>+ H1-TMD-H1 (5 $\mu\text{M}$ Rh-PE) |
| H1-TMD-H3 | SLB: POPC/cho/PEG-PE/NBD-PE (55/40/5/0.05 mol %)<br>+ H1-TMD-H1 (5 $\mu\text{M}$ Rh-PE) |

Images were acquired and analyzed by SymphoTime64 To obtain a fitted curve, the deconvolution method with calculated IRF was performed. All the fitting were the same as described in FLIM.

### Plant material and growth conditions

Seeds of *Nicotiana benthamiana* wild type (WT) and FX-KO plants (Ref) were germinated on peat pellets and watered using modified Knopp’s nutrient solution. Growing conditions were set to a long day photoperiod (16 h of light, 8 h of darkness, 25°C) at a light intensity of 150 μmol of photons m-2 s-1 in a climate-controlled phytotron chamber. After four weeks, plants were transferred to soil and grown for an additional two weeks. Six-week-old plants were used for transformation.

### Construct design

The native H1 (SKJ4) and mutated H1 (SKJ5) coding sequences (CDS) of the HA protein were codon-optimized for expression in *Nicotiana benthamiana* and used for subsequent subcloning into a tobacco-optimized CPMV RNA pEAQ-HT vector (Sainsbury et al., 2009) using type IIS restriction enzyme digest (BshTI and XhoI) and ligation (T4). Affinity purification tags (10xHis and TAP) were placed downstream of the CDS of the HA genes. Factor Xa protease cleavage site was included to optimize cleavage in vitro at HA1/HA2 site. Constructs were sequenced for quality control and propagated in *Escherichia coli* for downstream transformation using standard molecular biology protocols.

### Agrobacterium and tobacco transformation

Electrocompetent *Agrobacterium tumefaciens* GV3101 (Agrobacterium) cells were transformed using a standard electroporation protocol and selected on YEB plates (0.5 % (w/v) beef extract, 0.1 % (w/v) yeast extract, 0.5 % (w/v) peptone, 0.5 % (w/v) sucrose and 0.05 % MgCl_2_) containing rifampicin (100 mg/L), gentamicin (40 mg/L) and kanamycin (50 mg/L). Positive colonies were streaked and cultured according to (Leuzinger et al., 2013). Six-week-old tobacco plants were used for transient transformation via agroinfiltration of whole plants. Plants were submerged completely in the infiltration buffer (10 mM MES, pH 5,5; 10 mM MgSO_4_) containing suspended Agrobacterium cells and 0.1% Triton X-100. Submerged upside down-placed plants were placed in a vacuum chamber and treated with a vacuum of 400 mbar for 60 seconds, after which the valve was released slowly to allow the penetration of Agrobacterium into interstitial spaces of plant leaves. This procedure was repeated twice for every plant. Vacuum-infiltrated plants were returned to the phytotron chamber and grown for an additional 7 days.

### VLP isolation

Plant leaf tissue was weighed and homogenized in the isolation buffer (0.1 M sodium phosphate, pH 7.2) supplemented with a protease inhibitor cocktail (cOmplete, Roche) using a Waring blender. The homogenate was filtered and centrifuged to remove cell debris at 15,000 x g for 20 min in 4°C. The extract was used for ultracentrifugation on a step sucrose gradient of 25-70% for 4 h at 275,000 x g in 4°C in a Beckman Coulter Optima XPN-100 ultracentrifuge. Isolated VLPs were dialyzed and concentrated for downstream experiments.

### Protein isolation and purification

Plant tissue was ground in liquid nitrogen, then transferred to the protein extraction buffer (50 mM Tris-HCl pH 8.0, 10% glycerol, 2% SDS, 25 mM EDTA, 1 mM PMSF). Suspended samples underwent three freeze-thaw cycles in the presence of ultrasounds. Lysates were centrifuged at 10,000 x g at 4°C to remove the cell debris. Protein concentration was determined, and supernatants were used for downstream analysis. Affinity chromatography was performed using tested laboratory protocols using NiNTA or IgG resins according to the manufacturer’s instructions.

### SDS-PAGE and Western Blot

SDS-PAGE and Western Blot immunodetection were performed using routine protocols. Immunodetection was visualized using a ChemiDoc MP scanner (Bio-rad). Antibodies against HA protein were obtained commercially.

### Proteoliposome reconstitution

Purified membrane proteins were reconstituted into LUVs of known lipid composition (64 and 221) by incubation of the protein with the LUVs in the presence of 1 % CHAPS for 15 minutes in 30°C at 600 rpm after which the samples were dialyzed in the excess of HBS buffer to remove the detergents. After dialysis, reconstituted proteoliposomes were separated from LUVs and the free protein via floatation assay ultracentrifugation on a sucrose step gradient (0-35%). Reconstituted proteoliposomes were dialyzed against the HBS buffer once more to remove the sucrose.

### Transmission Electron Microscopy

To confirm the self-assembly of HA-VLPs in planta, transmission electron microscopy (TEM) imaging of transformed tobacco leaves and VLP isolates was performed. Sample preparation was performed as previously described (Kowalewska et al., 2016). For imaging we used JEM 1400 electron microscope.

