## Supplementary Information for "Transmembrane domains of H1 and H3 hemagglutinin contribute to membrane fusion in a different manner"

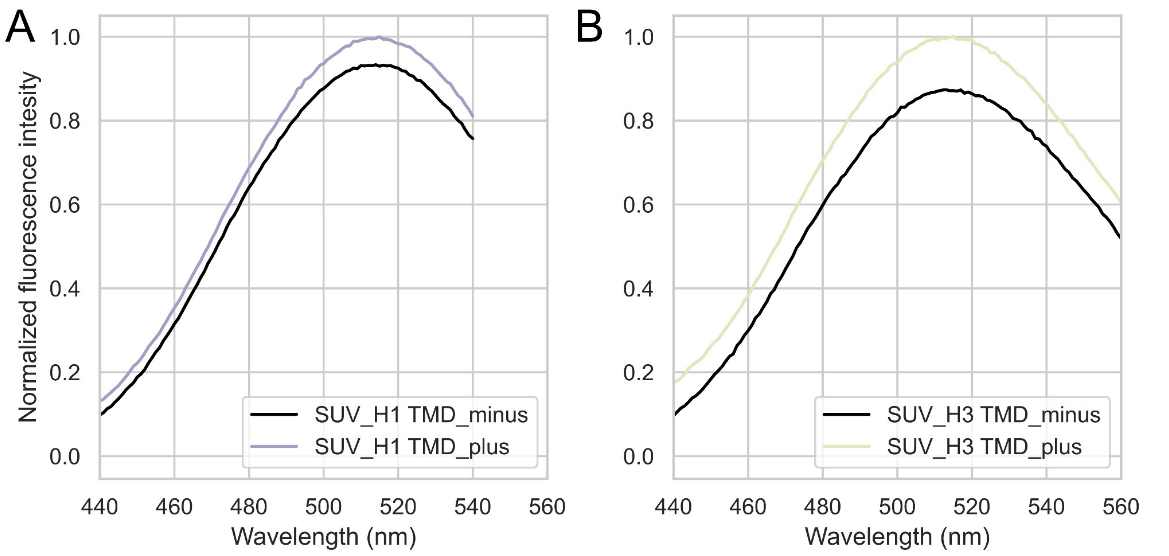


**SI Fig. 1** Incorporation of peptides into bilayers: **(A)** FRET between tyrosine (from H1 TMD) and Dansyl-PE, **(B)** FRET between tryptophan (from H3 TMD) and Dansyl-PE, SUV composition: POPC with 1 mol% Dansyl-PE, pH = 5.

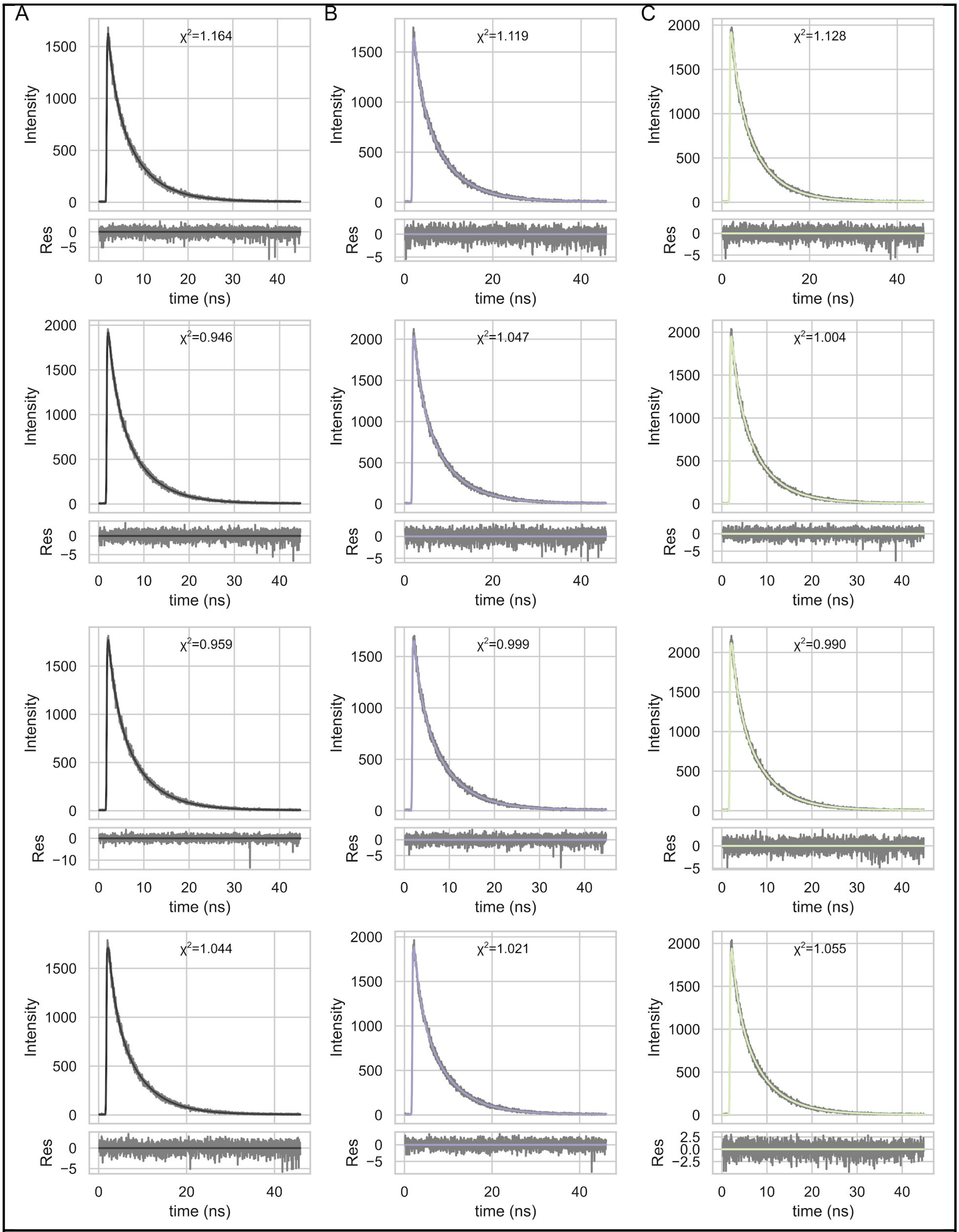


**SI Fig. 2** Examples of two-exponential tail fits of NBD-C6-PE fluorescence decays. Left pannel: pure POPC membrand, middle pannel: with H1 TMD peptide, right pannel: with H3 TMD peptide.

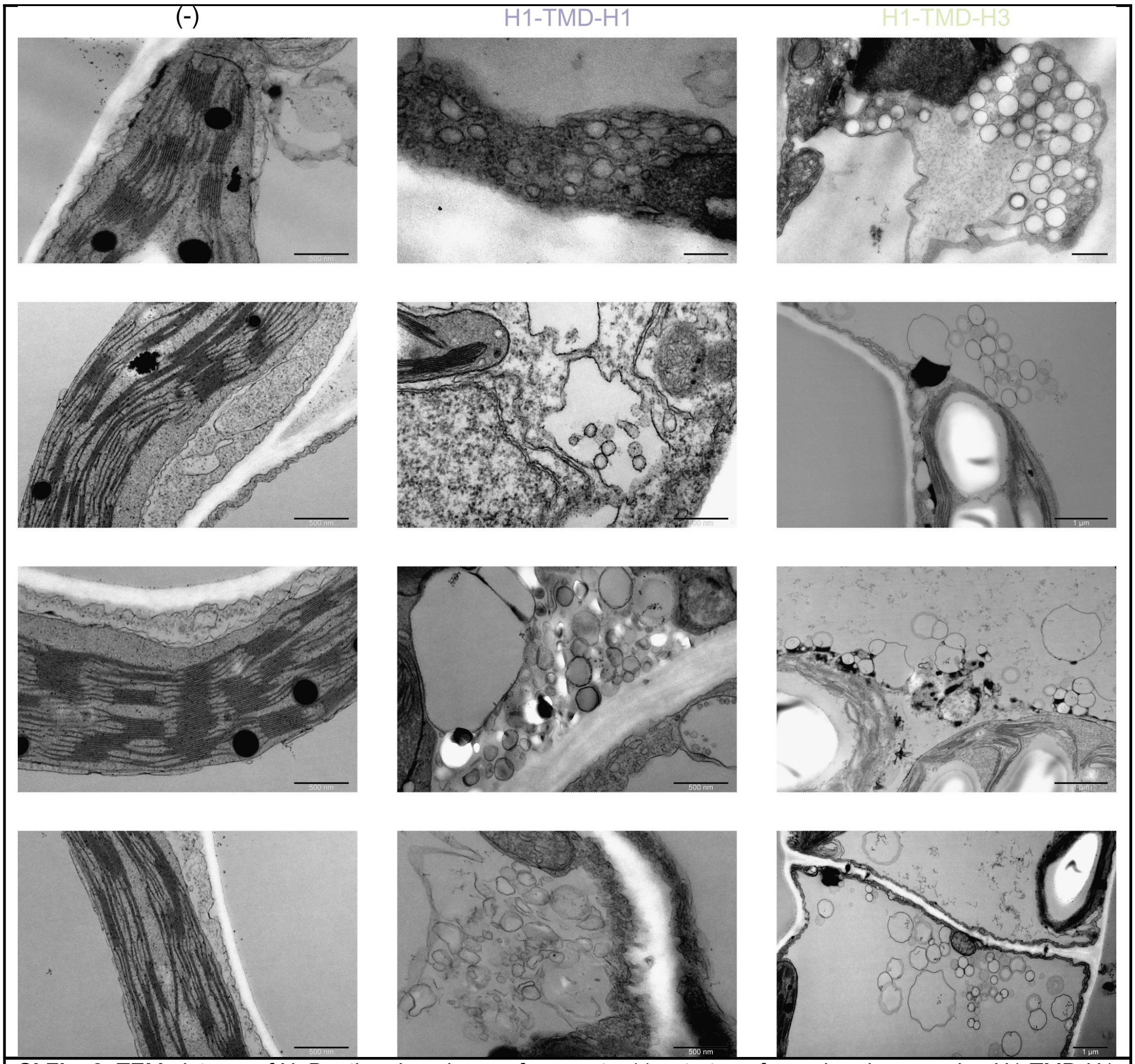


**SI Fig. 3** TEM pictures of N. Benthamiana leaves fragments: (-) not ntransformed and expressing H1-TMD-H1 and H1-TMD-H3 constructs (middle and right pannels, respectively).

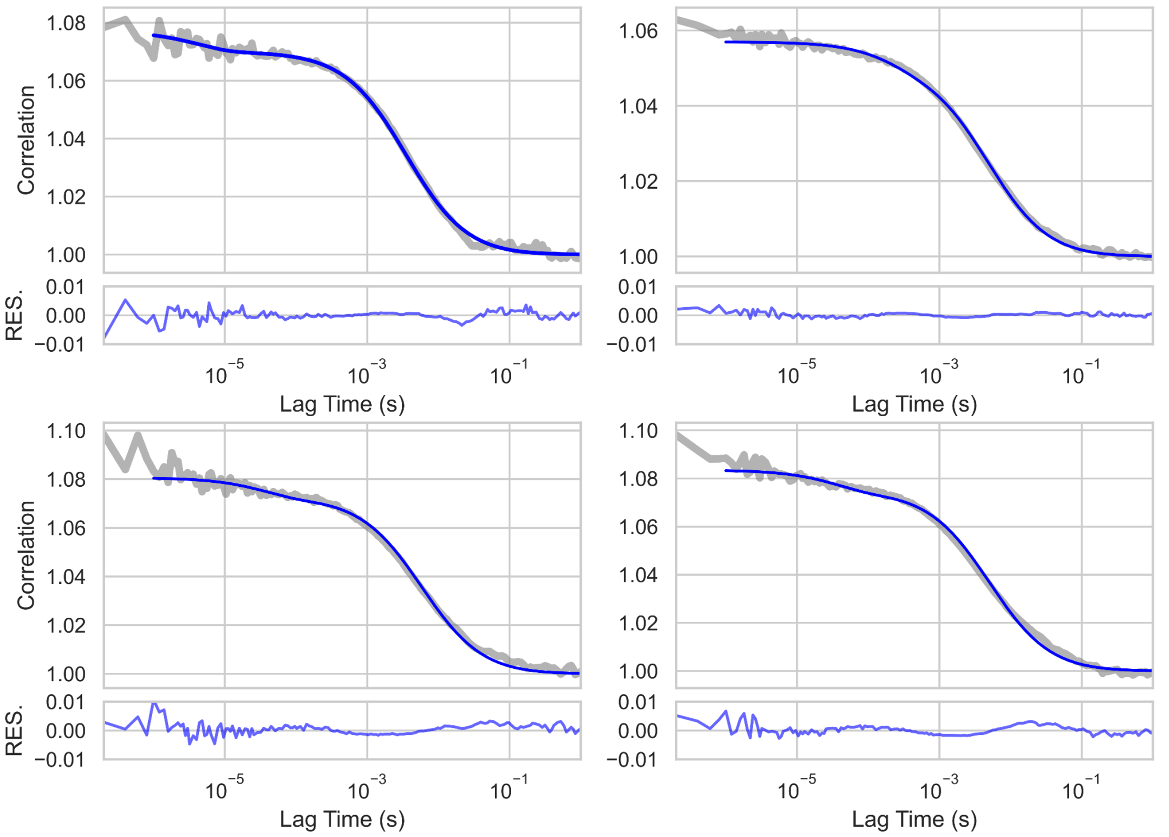


**SI Fig. 4** Mobility of lipids in SLB (POPC/chol/PEG-PE/Atto 488 (55/40/5/0.02 mol %)) examplified by FCS autocorrelation curves (55/40/5/0.02 mol %). The grey line represents the autocorrelation curve, the blue line shows the fitted curve with corresponding residuals plots. The diffusion coefficient obtained from the experiments is D=1,82 ± 0,61 $\mu m^2/s$, n=19.

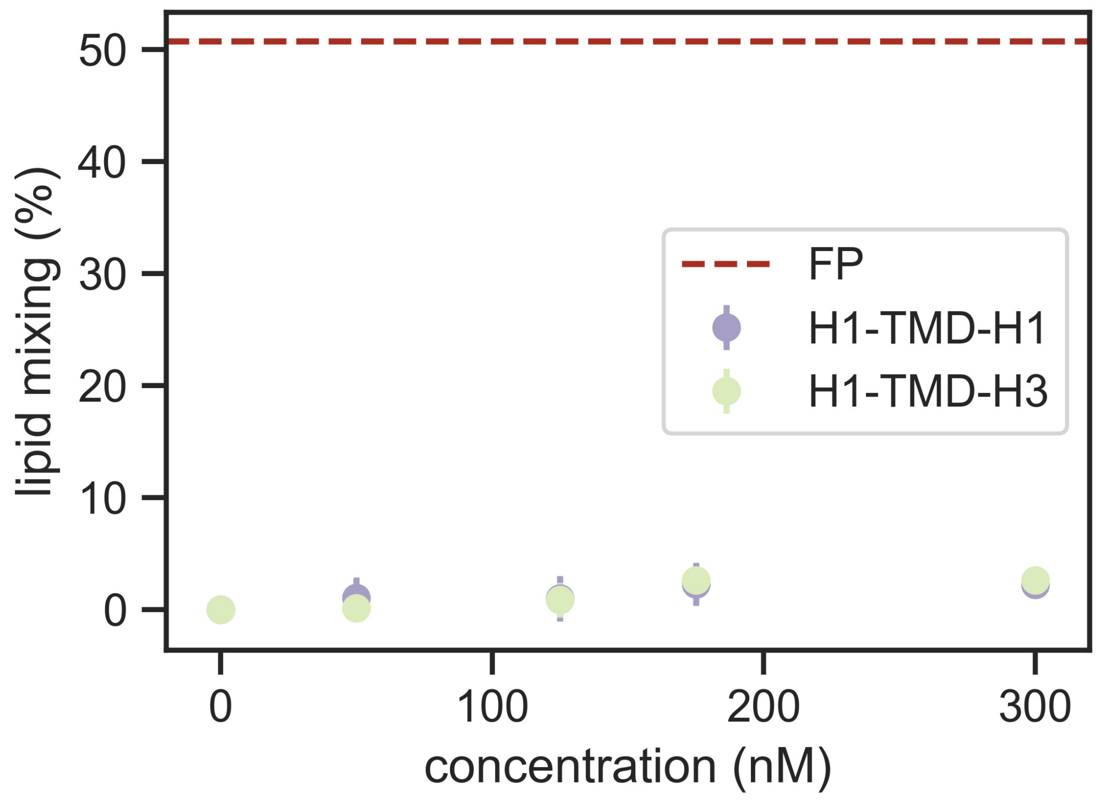


**SI Fig. 5** Lipid mixing experiment performed using cleaved HA in VLPs in pH 7.0.

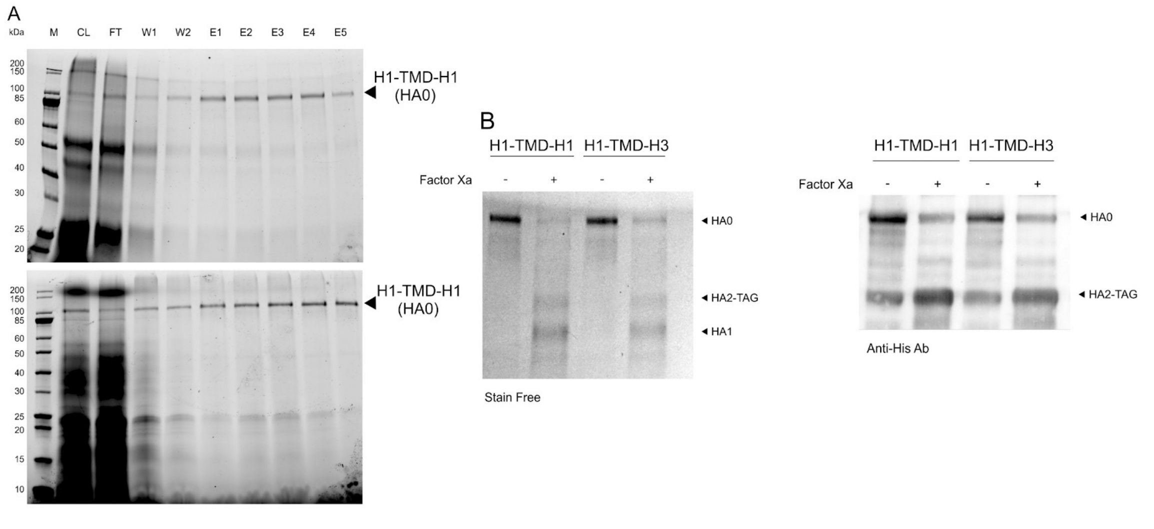


**SI Fig. 6 (A)** HA purification fractions (M- marker, CL- column load, FT- flow through, W1-W2 – low concentration imidazole wash, E1-E5 elution fractions. **(B)** Cleavage of purified proteins (left pannel), a Western blot using Anti-His antibody – right pannel.

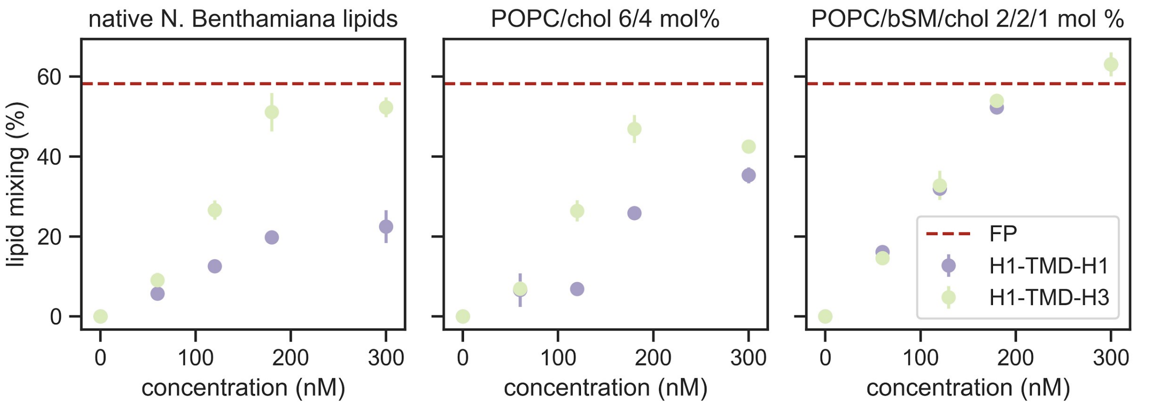


**SI Fig. 7** Lipid mixing experiment performed using HA in native *N. Benthamiana* VLPs (left pannel) and reconstituted in LUV of a known composition of choice (POPC/cholesterol 6/4 mol% - middle pannel) and POPC/brain sphingomuelin/cholesterol 2/2/1/ mol% - right pannel).
